# A Temporal Reliability and Sustainability Score Framework for Ranking Heart Rate Variability Metrics During Pharmacological Autonomic Blockade in Rats

**DOI:** 10.64898/2026.09.04.748642

**Authors:** M. Khawar Ali, Shuangshuang Ji, Siden Long, Shiyuan Gong, Jiande D.Z. Chen

**Affiliations:** Division of Gastroenterology and Hepatology, Department of Internal Medicine, School of Medicine, University of Michigan, Ann Arbor, MI, USA

## Abstract

**Background:** While Heart Rate Variability (HRV) is a cornerstone of non-invasive autonomic assessment, the sensitivity and sustainability of specific HRV parameters remain debated. This study introduces a “sustainability score” to systematically validate which metrics best reflect autonomic nervous system (ANS) modulation following pharmacological blockade.

**Methods:** Ten male Sprague-Dawley rats were implanted with ECG electrodes. After recovery, rats underwent randomized atropine (1 mg/kg, intraperitoneal) and propranolol (10 mg/kg, intraperitoneal) sessions separated by at least 3 days. Baseline ECG was recorded for 30 min before each session, followed by 90-min post-drug ECG recording. HRV parameters were analyzed in sequential 15-min and 30-min segments. Reliability was assessed using average p-values across post-drug segments and a sustainability score, defined as the number of segments significantly different from baseline.

**Results:** After atropine, HF Power showed the strongest rapid parasympathetic-related response in 15-min segments, with the lowest average p-value (0.1476) and highest sustainability score (4), followed by RSA (p=0.16148; score=4). HFn was less reliable in 15-min segments (p=0.23147; score=2) but became the most reliable parasympathetic-related metric in 30-min segments (p=0.0001; score=3). After propranolol, SI was the most reliable sympathetic-related metric in both 15-min (p=0.18225; score=2) and 30-min segments (p=0.0393; score=2), while SD2/SD1 showed the strongest 30-min sympathovagal-balance response (p=0.0200; score=3).

**Conclusion:** HF Power is recommended for rapid parasympathetic-related assessment in short windows, HFn for longer-window parasympathetic assessment, SI for sympathetic-assessment, and SD2/SD1 for 30-min sympathovagal-balance analysis. Sustainability scoring provides a structured framework for selecting ECG-derived autonomic markers.

## Introduction

Heart Rate Variability (HRV) reflects the dynamic effect of sympathetic and parasympathetic branches of the autonomic nervous system (ANS). Several mathematical approaches in time and frequency domains as well as nonlinear techniques have been employed to capture the complexity of the autonomic regulation; however, they may not exclusively represent a single autonomic branch. Identifying the most reliable HRV parameters for reflecting sympathetic and parasympathetic activity is crucial for advancing our understanding of autonomic function and for developing therapeutic strategies targeting autonomic dysfunction in various diseases like Functional Gastrointestinal Disorders (FGIDs)(Rajendra Acharya et al. 2006; Ali and Chen 2023). To distinguish between the sympathetic and parasympathetic contributions to the heart rate variability, the use of pharmacological interventions has been used in this study. Pharmacological agents can significantly alter the HRV metrics, but the specific temporal dynamics and the most responsive parameters have not been fully elucidated.

The autonomic function can be manipulated by using pharmacological agents like propranolol which blocks sympathetic activity and atropine which blocks the parasympathetic activity. Propranolol is a non-selective β-adrenergic receptor antagonist which is widely used to block the sympathetic activity by blocking the action of the endogenous catecholamines, epinephrine and norepinephrine at beta adrenergic receptors. Administering propranolol leads to the reduction in heart rate and a decrease in HRV parameters associated with sympathetic tone (Bylund 2015; Ammar and Hussein 2018).

Atropine is a non-selective muscarinic receptor inhibitor, which inhibits parasympathetic activity by blocking the vagal influence on the heart. Muscarinic receptors are receptor sites for neurotransmitters of the parasympathetic nervous system (PNS), acetylcholine (Ach), located on the post-synaptic cell membranes of smooth muscle, cardiac muscle and glandular tissue at the end of the PNS. Thus, the muscarinic receptors mediate the physiological effects of parasympathetic nerve activities like cardiac slowdown, smooth muscle contraction, vasodilation and increased secretion of exocrine glands. Therefore, the atropine increases in heart rate and causes a reduction in parasympathetic-driven HRV parameters (Gordan, Gwathmey, and Xie 2015; Scott and Fryer 2012; Perera et al. 2017; Broadley and Kelly 2001). By modulating the autonomic nervous system selectively, these agents enable us to assess the relative contributions of sympathetic and parasympathetic activity to the HRV and validate HRV parameters.

The aim of this study was to validate physiological implications of HRV by systematically evaluating changes in various HRV parameters with the use of pharmacological blockade of sympathetic and parasympathetic activity in normal rats. The sensitivities of several commonly used HRV parameters in responses to these two agents were assessed in multiple experimental sessions. By analyzing the temporal changes in HRV parameters following the administration of these agents, we aimed to identify the most responsive and reliable HRV parameters that accurately reflected sympathetic and parasympathetic modulation.

## Methods

### Animals

Ten Adult male Sprague-Dawley (SD) rats, weighting 250–300g were used in this study. The rats were housed plastic cages placed in rooms with temperature of 22–23°C, humidity of 40–50%, and on a12:12-h light-dark cycle. Water and food were available ad libitum. The study protocol was approved by the Institutional Animal Care and Use Committee.

### ECG Electrodes Implantation

After one week of acclimation, the animals were subject to the following procedure. Under anesthesia with inhalation of 1.5–2.0% isoflurane, the rats were implanted with three subcutaneous electrodes used to record the electrocardiogram (ECG). After disinfection, the hairs were shaved, the skin on top of the corresponding position was cut open, three chest skin incisions (about 2 mm) were made to implant three cardiac pacing wire (Medtronic, Minneapolis, MN) electrodes on the left and right chest subcutaneous area and on the left side of the cardiac apex subcutaneous, respectively. The lead wires were tunneled underneath the skin and externalized percutaneously on the rat’s neck as shown in figure 1 below (adapted from (Wang et al. 2019). After the surgical procedure, Carprofen (5 mg/kg) and antibiotic drug enrofloxacin (5 mg/kg) were injected subcutaneously, daily for 3 days starting from the day of surgery to provide postoperative pain relief. The rats were housed in individual cages to protect the electrodes wires from being chewed off by other rats. The experiment was conducted after one week recovery period post the surgical implantation of electrodes.

**Figure 1:**
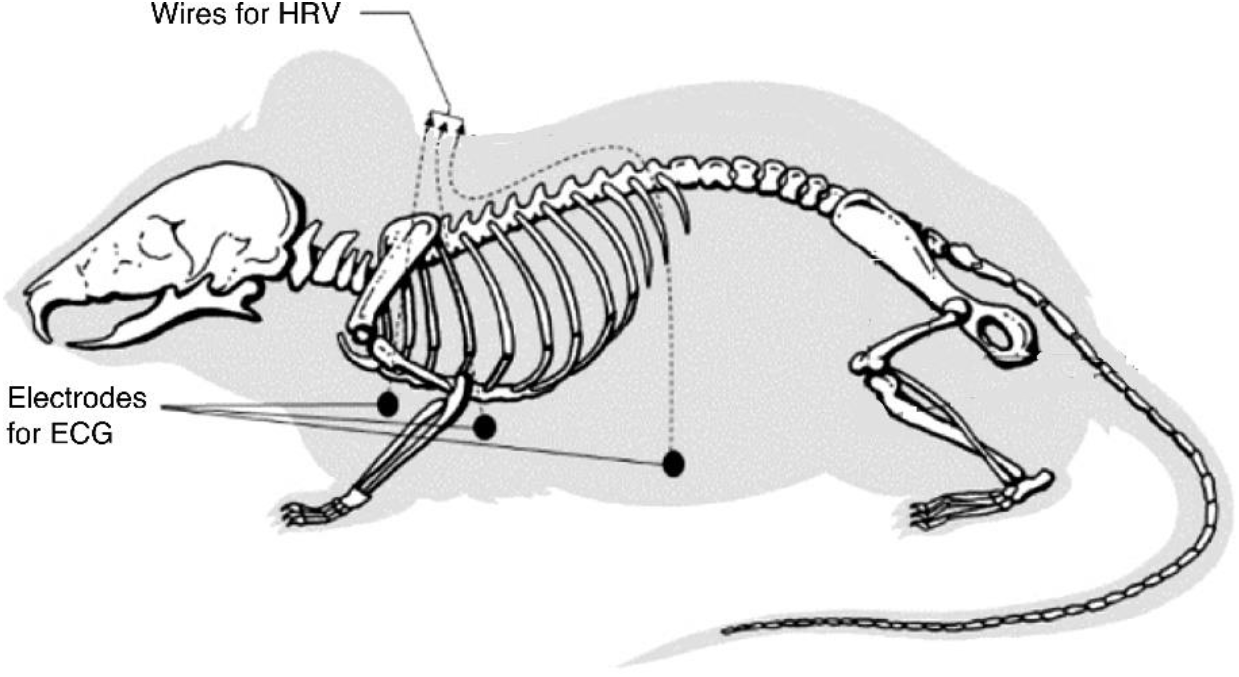
ECG electrodes implantation in rats, figure adapted from (Wang et al. 2019)

### Experimental Protocol

After recovery from the surgical implantation of ECG electrodes for 7 days, rats were fasted overnight before the day of experiment. On the day of experiment, the animal was placed in a transparent strainer, and the baseline ECG was recorded for 30 min using a special amplifier (model 2283 Fti Universal Fetrode Amplifier, UFI, Morro Bay, CA, USA).

All 10 rats completed the propranolol experiment. Three animals did not complete the atropine experiment because of mortality before the atropine session; therefore, atropine analyses were performed in the remaining seven animals. The cause of mortality was not determined, and no additional mortality occurred during atropine recordings.

After baseline recording, rats underwent two separate pharmacological blockade sessions in randomized order, separated by a minimum 3-day washout period. In one session, atropine was administered intraperitoneally at 1 mg/kg to block muscarinic parasympathetic activity. In the other session, propranolol was administered intraperitoneally at 10 mg/kg to block β-adrenergic sympathetic activity. A new 30-min baseline ECG recording was obtained before each drug administration. After injection, rats were returned to the restrainer, and ECG was recorded for 90 min. Some rats received atropine first, whereas others received propranolol first.

MATLAB code developed and validated by the first author was used for HRV analysis, which has already been used in earlier studies (Ali et al. 2023; 2021; Yuan et al. 2020)). The code was developed to remove the artifacts from the recorded ECG signal, detect the R-peaks from the ECG signal, generate the RR interval signal (HRV signal) and to calculate the HRV parameters from the RR interval signal. The HRV parameters were calculated for 30-min baseline recording in each case, and for 15 min as well as 30 min segments after the drug administration to study the sensitivity and robustness of the HRV parameters.

Sympathetic Index (SI) which has been used as a pure measure of sympathetic activity (Baevsky and Chernikova 2017; Ali et al. 2023; 2021; Yuan et al. 2020), Low Frequency (LF) power and normalized Low frequency (LFn) which are widely used as marker of sympathetic activity but are known to be influenced by both the sympathetic as well as parasympathetic activities (Shaffer and Ginsberg 2017; Ali et al. 2023), and the major axis of Poincare plot (SD2) were tested against the sympathetic blockade via propranolol. LF/HF ratio and Poincare plot major axis to minor axis ratio (SD2/SD1) were tested against propranolol as marker of the autonomic balance. Similarly, the parasympathetic parameters of heart rate variability, root mean square of successive differences of RR interval (RMSSD), high frequency power band (HF Power), normalized HF Power (HF/(HF+LF)), Respiratory Sinus Arrhythmia (RSA), minor axis of the fitted ellipse of the Poincare plot (SD1), pNN50%, SDSD and SDNN were tested in response to the parasympathetic blockade via atropine in 15 minutes as well as 30 minutes segments compared to the baseline (Ali et al. 2023; Shaffer and Ginsberg 2017; Ali et al. 2021).

Some modifications were made to the calculation procedures of the HRV parameter formulae generally used in human studies to better suit the analysis in animal studies. For calculating Sympathetic Index (SI), the bin width of the histogram was changed to 15ms instead of 50ms used in human studies due to very high heart rate which corresponds to very small inter-beat intervals in rats compared to the humans (Ali et al. 2021; Yuan et al. 2020; Ali et al. 2023). Similarly, the power band for HF power was taken from 0.8 to 3.0 Hz. reflecting cardiac vagal efferent activity and for LF, it was taken from 0.2 to 0.8 Hz. (Yin, Chen, and Chen 2010).

### Statistical Analysis

The HRV parameters were calculated for the entire 30-min baseline recording and in 15-min segments for the 90-min after the drug administration (1-15 min, 16-30 min, 31-45 min, 46-60 min, 61-75 min and 75-90 min) as well as in 30-min segments for 90 minutes after the drug administration (1-30 min, 31-60 min and 61-90 min). The comparison was carried out between the baseline value vs all the 15 min segments followed by the comparison between the baseline and all 30-min segments post drug administration to study the sensitivity of each HRV parameter. Shapiro-Wilk normality test was used to test for normal distribution of data before conducting the comparison. If the data was normally distributed, one-way ANOVA followed by Holm-Sidak multiple comparison test was applied. While for data which was not normally distributed, Friedman test followed by Dunn’s multiple comparison test was used for comparison. The significance level was set for p<0.05. All the statistical analyses were carried out in GraphPad Prism software.

Finally, the p-values of all the HRV variables were compiled across all the 15-minute and 30 minutes segments after the drug administration and their average p-value was calculated. A sustainability score which is equal to the number of segments in each case (15 min and 30 min), the HRV variable remained significantly different compared to its baseline value after the drug administration was calculated. The variable with the highest sustainability score and the lowest average of its p-values was selected as the most reliable parameter.

## Results

### Validation of HRV parameters reflecting parasympathetic activity

Atropine is known to act within 10 to 15 minutes after the IM administration, with a duration of action of 90 minutes (“Intramuscular Sedation” 2018). Blocking parasympathetic activity via atropine injection caused an increase (p=ns) in the average heart rate within the first 15 minutes which continued to diminish until the 90 minutes, As shown in the figure 2 below, the average value of HR increased by 11.81% in the first 15 minutes (p=ns) and then started to decrease and kept on decreasing during the following 15-minutes segments, it went below the baseline value in 45-60min (0.89% below the baseline), 60-75 min (4.38% lower than the baseline value) and 75-90 min (6.11% lower than the baseline value). However, none of these changes attained statistical significance (p=ns).

**Figure 2:**
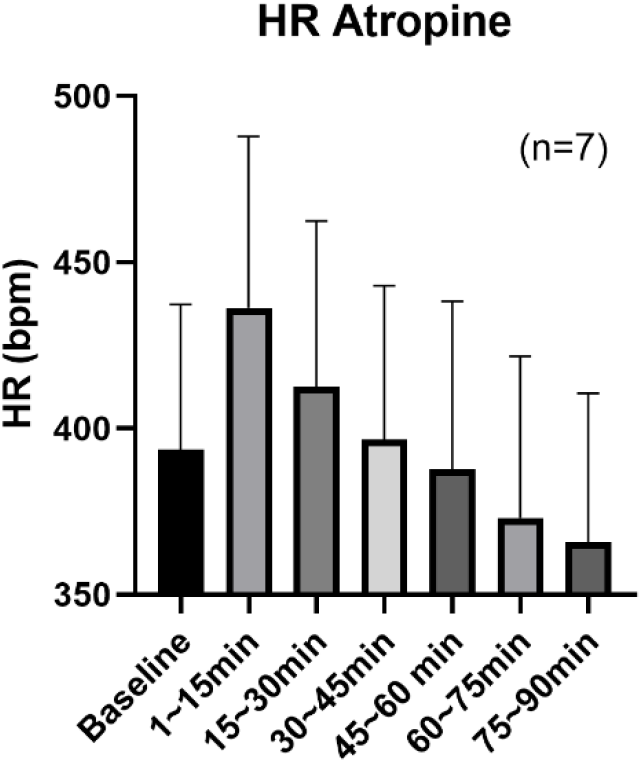
Heart Rate modulation by Atropine administration in rats

RMSSD did not show significant change within the first 15 minutes of atropine administration although its average value was suppressed from 16.54±17.33ms during baseline to 1.62±0.56ms during the first 15 minutes (p=ns). However, a significant decrease was observed in RMSSD, for 15-30 minutes (p=0.0002) and 30-45 minutes (p=0.0078) compared to its baseline value. After 45 minutes, the values of RMSSD started to recover, and no statistical difference was observed compared to its baseline value (Figure 3A).

**Figure 3:**
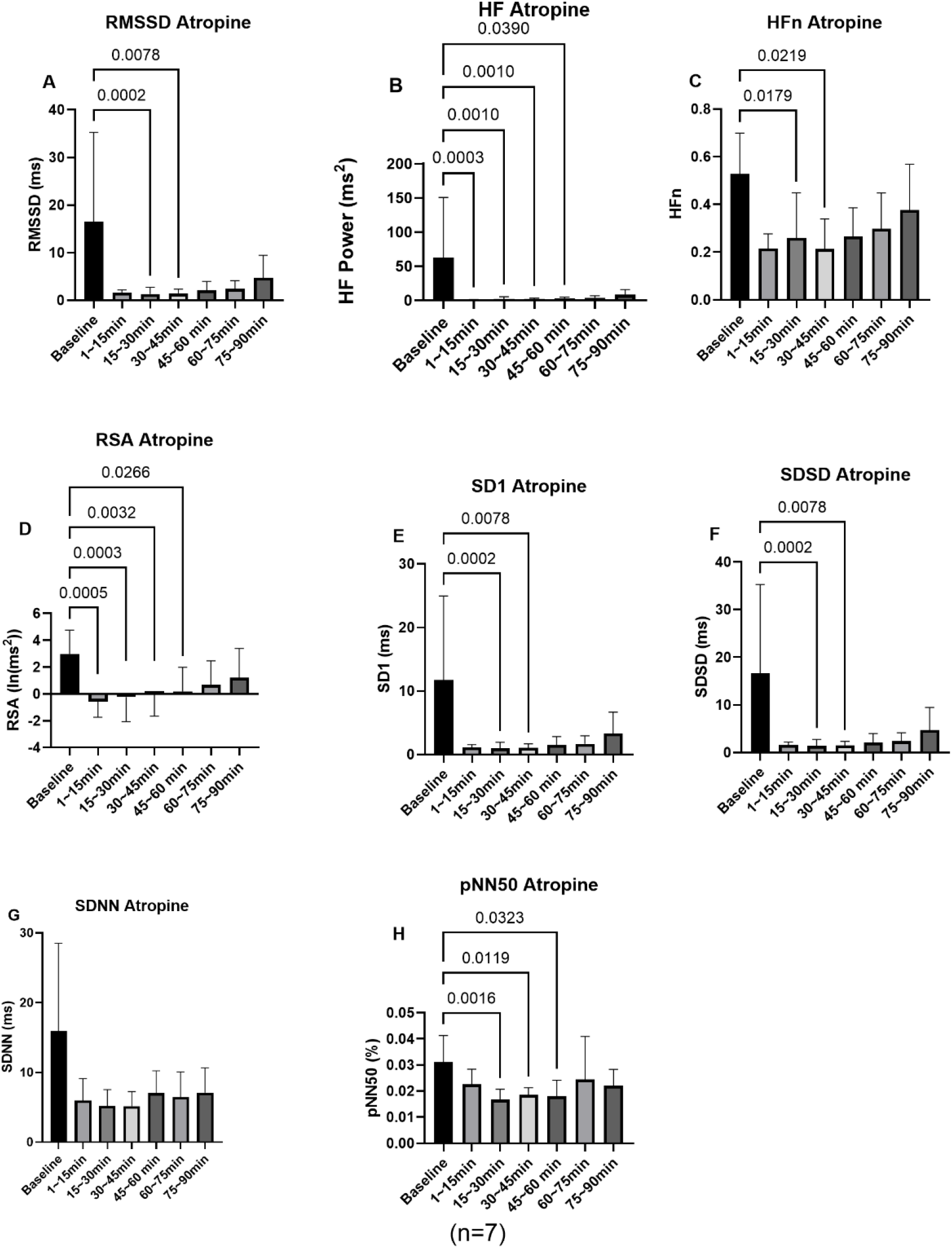
Comparison of HRV parameter values in segments of 15-mins after the IP administration of Atropine in 7 rats (N=7) up to 90 minutes with their respective baseline values (A)RMSSD, (B) HF Power (C) HFn (D) RSA (E) SD1 (F) SDSD (G) SDNN (H)pNN50%

High Frequency power (HF power) and RSA (which is derived from the HF power) on the other hand indicated quick and sustained responses to atropine administration. HF power was significantly decreased in the first 15 minutes (p=0.0003) and stayed low for 15-30 minutes (p=0.0010), 30-45 minutes (p=0.0010) and 45-60 minutes (p=0.0390), however, no significant change was observed after 60 minutes (Figure 3B). Similarly, the RSA was significantly decreased in the first 15 minutes (p=0.0005) and stayed low for 15-30 minutes (p=0.0003), 30-45 minutes (p=0.0032) and 45-60 minutes (p=0.0266), and it started to recover after 60 minutes, when no significant change was observed (Figure 3D). While normalized HF power, which was calculated as a ratio of HF power to the sum of HF and LF powers, didn’t show significant change in the first 15 minutes (p=ns) compared to its baseline value but decreased for 15-30 minutes (p=0.0179) as well as for 30-45 minutes (p=0.0219). After 45 minutes, no change was observed (Figure 3C).

Both SD1 and SDSD were not decreased significantly (p=ns) in the first 15 minutes, while the significantly decreased parasympathetic activity was captured during the 15-30 minutes (SD1: p=0.0002, SDSD: p=0.0002) as well as 30-45 minutes (SD1: p=0.0078, SDSD: p=0.0078), while no change was observed after 45 minutes (Figures 3E and F). The response of SD1 and SDSD was exactly like that of RMSSD. SDNN on the other hand, didn’t change significantly with the atropine, although its average values decreased from 15.95±11.64 ms during baseline to 5.95±2.93ms in the first 15 minutes (p=ns) and stayed below 7.5ms throughout the 90 minutes (Figure 3G). Although the absolute values of pNN50%, were very low compared to the human studies (Ali et al. 2023), it was still able to capture the suppressed parasympathetic activity during 15-30 min (p=0.0016), 30-45 minutes (p=0.0119) and 45-60 minutes (p=0.0323), while no change was observed after 60 minutes (Figure 3H).

As shown in table 1 above, HF power had the lowest average of its p-values (p_ave=0.1476) calculated from all the 15-minute segments with the highest sustainability score of 4. While RSA has the second lowest average p-value of 0.16148 with same sustainability score as HF power. RMSSD, SD1 and SDSD had the same average of its p-values as well as the same sustainability score. pNN50% had sustainability score of 3 but highest average of it p-values across all 15-minute segments (Table 1, Figure 4).

**Table 1:** p-values and sustainability score for PNS variables.

|  | 1-15 min | 15-30 min | 30-45 min | 45-60 min | 60-75 min | 75-90 min | Average p-value | Sustainability Score |
| --- | --- | --- | --- | --- | --- | --- | --- | --- |
| HF Power | 0.0003* | 0.001* | 0.0010* | 0.039* | 0.1124 | 0.7319 | 0.1476 | 4 |
| RSA | 0.0005* | 0.0003* | 0.0032* | 0.0266* | 0.1124 | 0.8259 | 0.16148 | 4 |
| RMSSD | 0.1557 | 0.0002* | 0.0078* | 0.1124 | 0.8259 | 1 | 0.35033 | 2 |
| SD1 | 0.1557 | 0.0002* | 0.0078* | 0.1124 | 0.8259 | 1 | 0.35033 | 2 |
| SDSD | 0.1557 | 0.0002* | 0.0078* | 0.1124 | 0.8259 | 1 | 0.35033 | 2 |
| pNN50% | 1 | 0.0016* | 0.0119* | 0.0323* | 0.5693 | 0.6466 | 0.37695 | 3 |
| HF<sub>n</sub> | 0.0562 | 0.0179* | 0.0219* | 0.0801 | 0.2127 | 1 | 0.23147 | 2 |
| SDNN | 0.2512 | 0.2512 | 0.2416 | 0.2512 | 0.2144 | 0.2512 | 0.24347 | 0 |

**Table 2:** p-values and sustainability scores for Parasympathetic Nervous system variables for 30-min segments analysis.

|  | 1-30 min | 30-60 min | 60-90 min | Average p-value | Sustainability Score |
| --- | --- | --- | --- | --- | --- |
| **HF<sub>n</sub>** | 0.0001 | 0.0001 | 0.0001 | 0.0001 | 3 |
| **HF Power** | 0.0002 | 0.012 | 0.0219 | 0.0114 | 3 |
| **RSA** | 0.0057 | 0.0138 | 0.0429 | 0.0208 | 3 |
| **RMSSD** | 0.0162 | 0.018 | 0.0384 | 0.0242 | 3 |
| **SD1** | 0.0163 | 0.018 | 0.0384 | 0.0242 | 3 |
| **SDSD** | 0.0162 | 0.018 | 0.0383 | 0.0242 | 3 |
| **SDNN** | 0.0376 | 0.0165 | 0.0279 | 0.0273 | 3 |
| **pNN50%** | 0.0981 | 0.2024 | 0.3604 | 0.2203 | 0 |

**Table 3:** p-values and sustainability scores for sympathetic-related and sympathovagal-balance HRV parameters in 30-min segments after propranolol administration.

|  | 1-30 min | 30-60 min | 60-90 min | Average p-value | Sustainability Score |
| --- | --- | --- | --- | --- | --- |
| **SI** | 0.0168 | 0.0281 | 0.073 | 0.0393 | 2 |
| **LFn** | 0.0118 | 0.2692 | 0.0466 | 0.1092 | 2 |
| **LF/HF** | 0.0168 | 1.0000 | 0.0281 | 0.3483 | 2 |
| **SD2/SD1** | 0.0400 | 0.0095 | 0.0105 | 0.0200 | 3 |
| **LF Power** | 1.0000 | 0.2498 | 0.2498 | 0.4999 | 0 |
| **SD2** | 1.0000 | 1.0000 | 1.0000 | 1.0000 | 0 |

**Figure 4:**
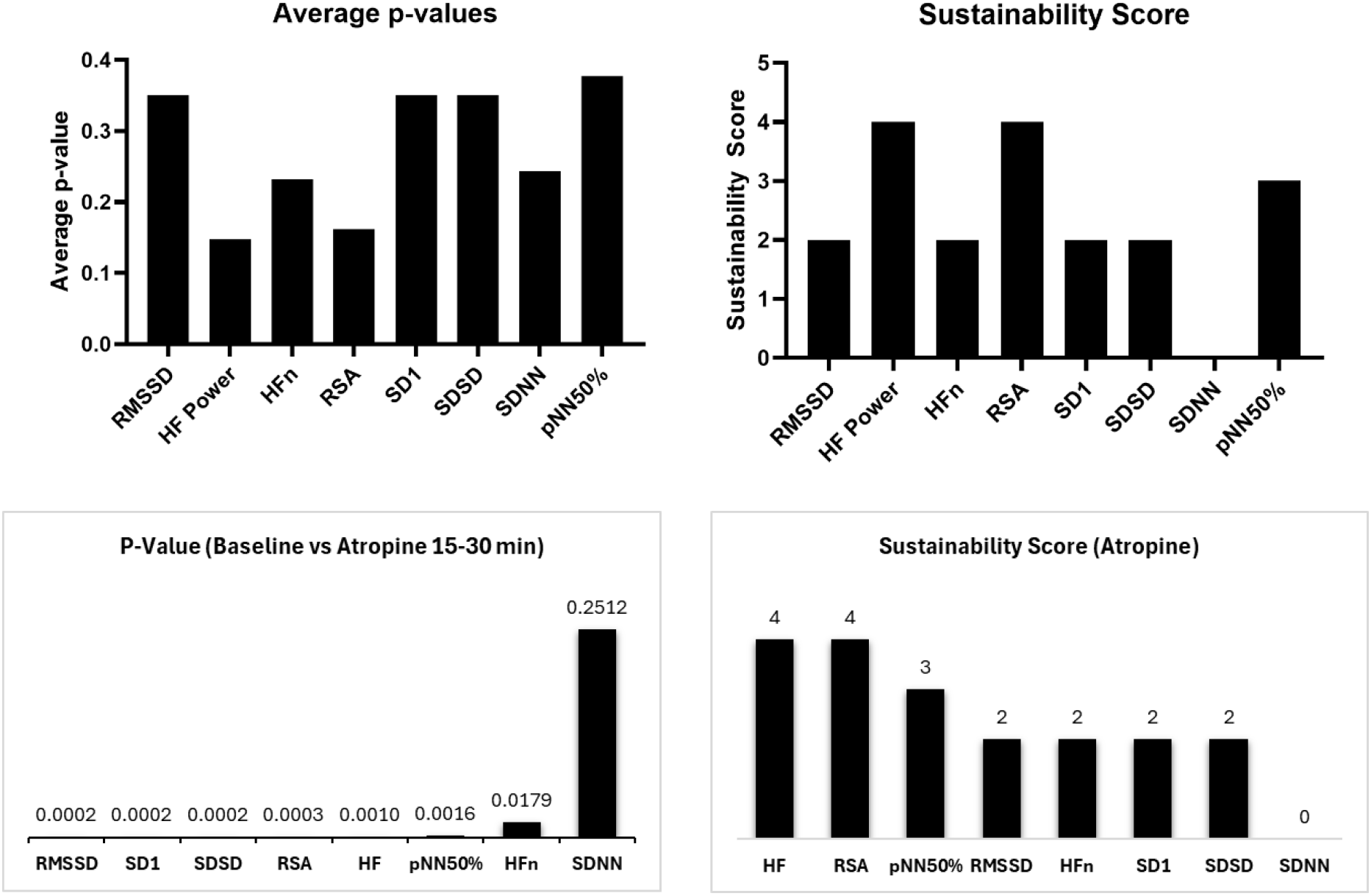
Summary ranking of parasympathetic-related HRV parameters after atropine administration. Average p-values were calculated across all 15-min post-atropine segments. Segment-specific p-values from the 15–30 min post-atropine window are shown in ascending order. Sustainability score was defined as the number of post-atropine segments significantly different from baseline.

### Validation of sympathetic-related and sympathovagal-balance HRV parameters after propranolol administration

Propranolol is known to act within 30-60 min and the effect lasts for 3-4 hours (PAterson et al. 1970; Coltart and Shand 1970).Blocking the sympathetic activity via propranolol caused a significant decrease in the heart rate after 15 minutes which continued to remain low till the 90^th^ minute, while the change in the first 15 minutes was not statistically significant although the average values decreased from 390.21±38.82 bpm during baseline to 295.39 bpm (suggesting a delayed effect of propranolol) as shown in the figure 5 below:

**Figure 5:**
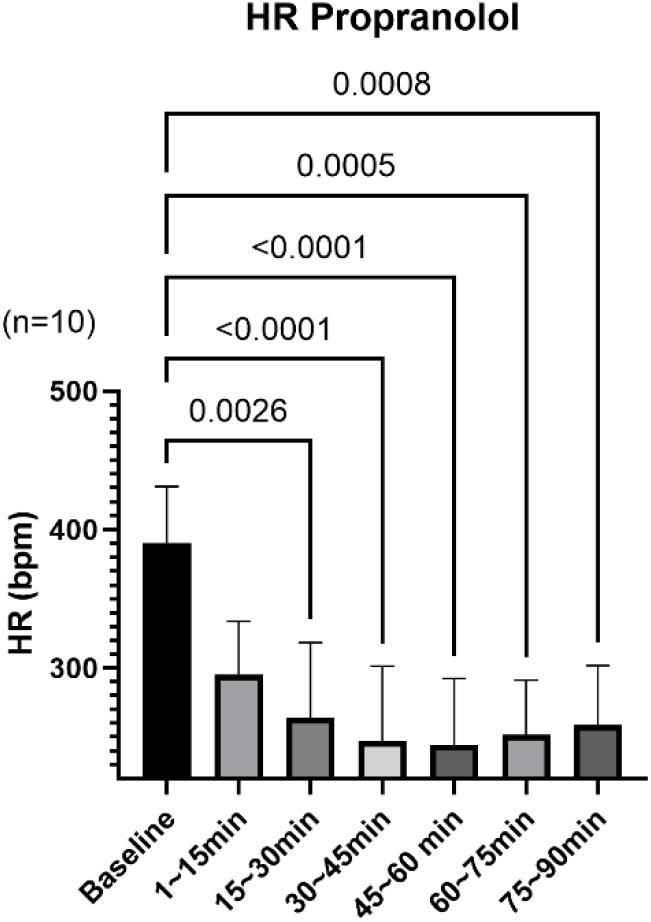
Heart Rate modulation by propranolol administration in rats

Sympathetic Index (SI) also did not show any significant change in the first 15 minutes after the administration of propranolol. The average value of SI decreased from 3444.46±3766.93 s^−2^ during baseline to 1178.01±867.89 s^−2^ during the first 15 minutes after the administration of propranolol, but, this change was not significant. However, for 15-30 minutes (p=0.0427) as well as 30-45 minutes (p=0.0026) the values of SI were significantly lower than that of baseline indicating the blocked sympathetic activity. While after 60 minutes, the SI started to recover back towards the baseline values, therefore, no significant difference was observed after 60 minutes as shown in figure 6A below.

**Figure 6:**
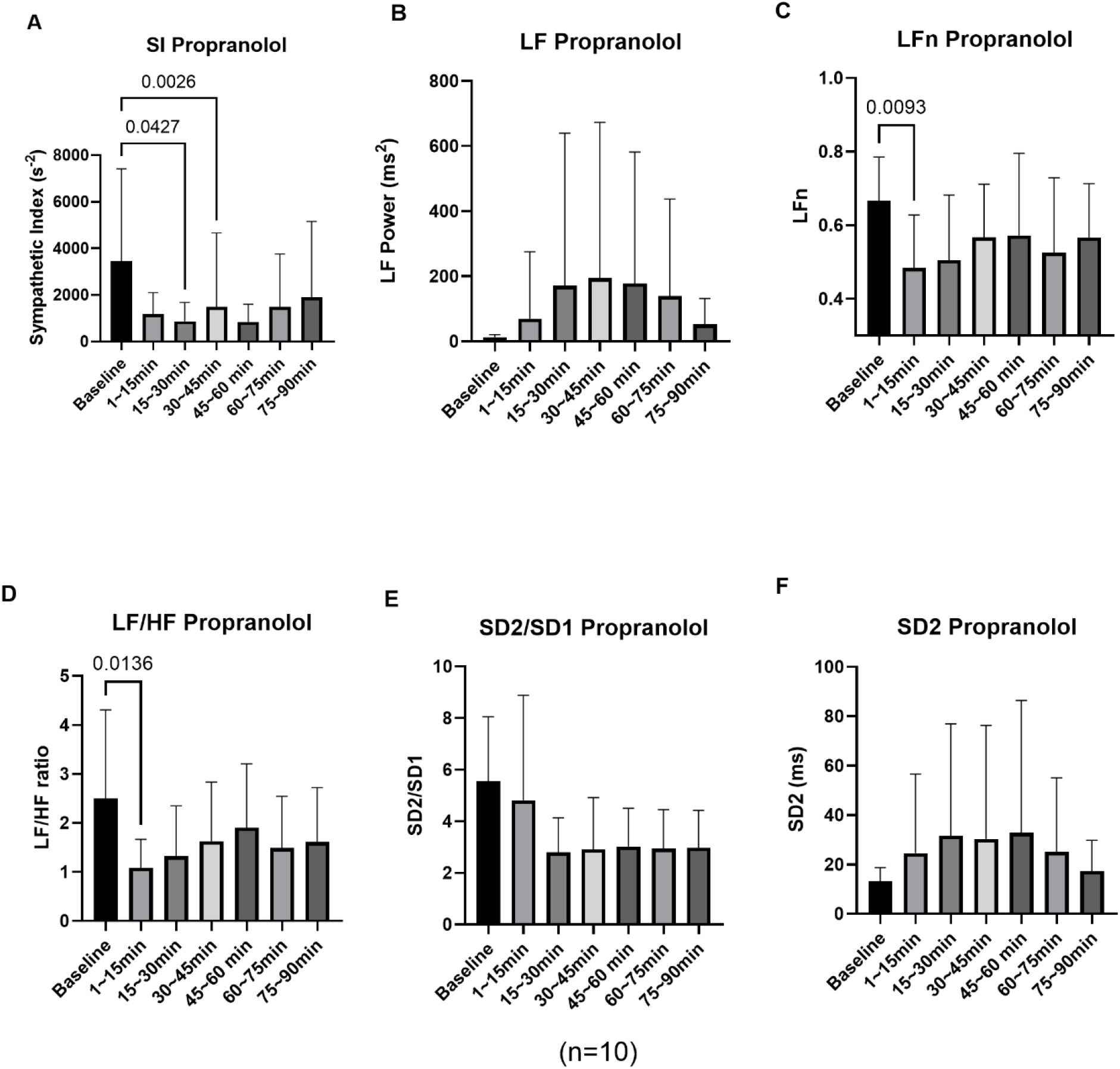
Comparison of HRV parameter values in segments of 15-mins after the IP administration of Propranolol in 10 rats (N=10) up to 90 minutes with their respective baseline values (A) Sympathetic Index (S), (B) LF Power (C) LFn (D) LF/HF Ratio (E) SD2/SD1 (F) SD2

**Figure 7:**
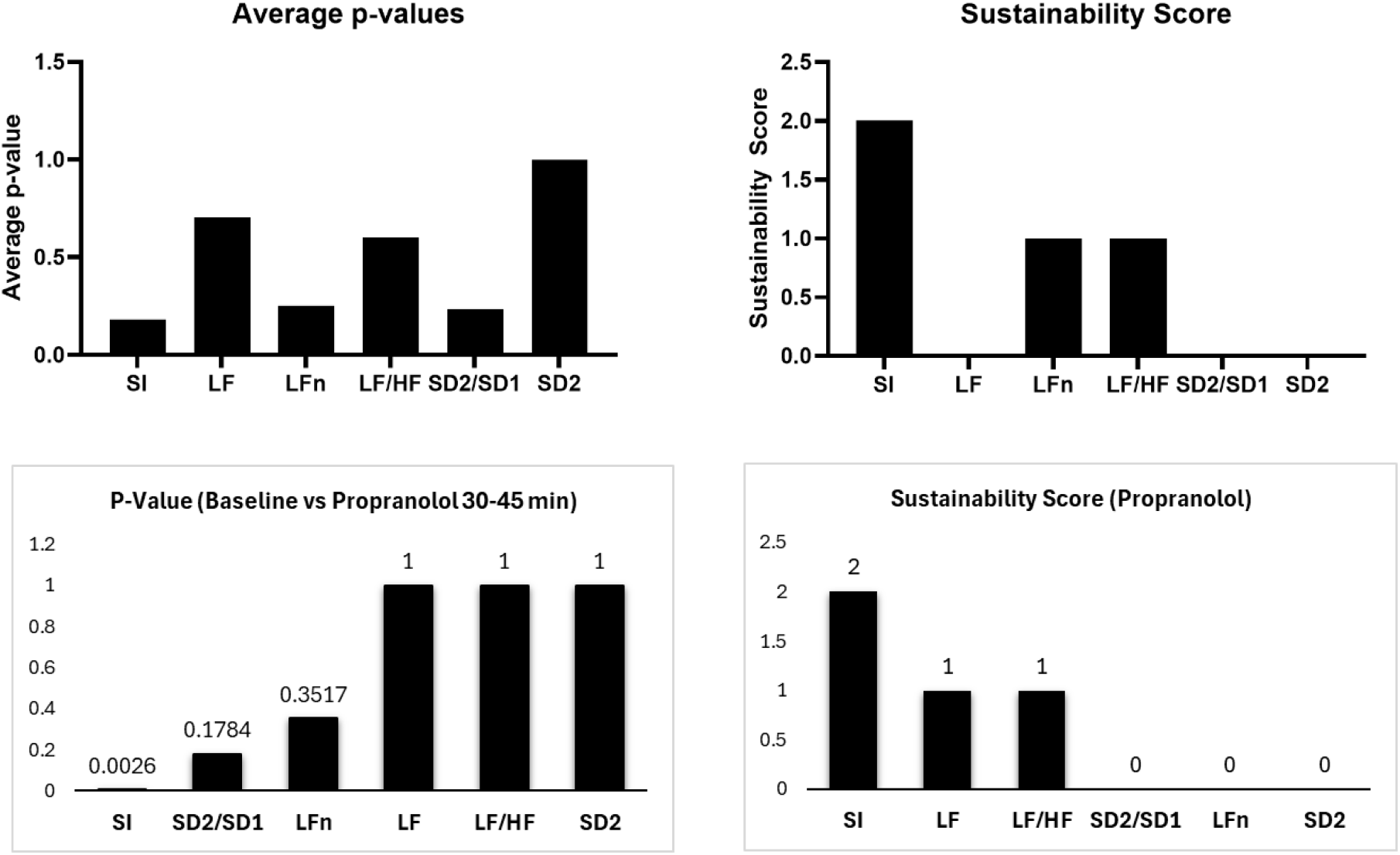
Average of p-values and sustainability score of sympathetic-related and sympathovagal-balance HRV parameters after propranolol administration

The low frequency (LF) power (Figure 6B) as well as the major axis of the Poincare plot (SD2) (Figure 6F)) didn’t show any change at all due to the administration of propranolol during any 15-min segment up to 90 minutes when compared to their respective baseline values. While normalized low frequency (LFn) power have shown significant change during the first 15 minutes only, no change was observed after 15 minutes compared to the baseline value (figure 6C).

On the other hand, the LF/HF ratio (Figure 6D), which is a marker of autonomic balance decreased significantly (p=0.0136) during the first 15 minutes, after 15 minutes the average values started to recover towards its baseline value with no significant difference observed. SD2/SD1 which also a marker of autonomic balance, did not show any significant change after the administration of propranolol during any 15-min segment up to 90-min compared to its baseline value (Figure 6E).

SI had the lowest average of its p-values (p-ave=0.18225) among all 15-minute segments post propranolol administration with the highest sustainability score of 2, followed by LFn with p-ave=0.2516, but its sustainability score was 1. SD2 had the highest average of its p-values with sustainability of zero. Among the parameters with Autonomic balance, LF/HF ratio had p-ave=0.5988 with a sustainability score of 1. SD2/SD1 was not significant during any 15-minute segment, hence, its sustainability score is 0, therefore, its average p-value is not considered.

### Effect of 30-min Analysis Windows on HRV Parameter Reliability

To determine whether HRV parameter reliability was influenced by recording duration, the post-drug ECG data were reanalyzed using non-overlapping 30-min segments. This analysis allowed comparison of parameter sensitivity and sustainability across longer recording windows relative to the 15-min segment analysis.

#### Parasympathetic-related HRV responses in 30-min segments after atropine administration

Among parasympathetic-related HRV parameters, most parameters detected parasympathetic suppression within the first 30-min segment and remained significantly different from baseline across the 1–30, 30–60, and 60–90 min post-atropine segments. However, pNN50% was not significant in any 30-min post-atropine segment, as shown in Figure 8. Notably, HFn showed improved reliability when HRV was analyzed using 30-min segments. While HFn showed only moderate reliability in the 15-min analysis, with an average p-value of 0.23147 and a sustainability score of 2, it had the lowest average p-value of 0.0001 and a sustainability score of 3 in the 30-min analysis. HF Power also remained highly reliable in the 30-min analysis, with the second lowest average p-value of 0.0114 and a sustainability score of 3. These findings suggest that HFn is more sensitive to parasympathetic blockade when longer HRV analysis windows are used, whereas HF Power shows strong reliability in both shorter and longer analysis windows.

**Figure 8:**
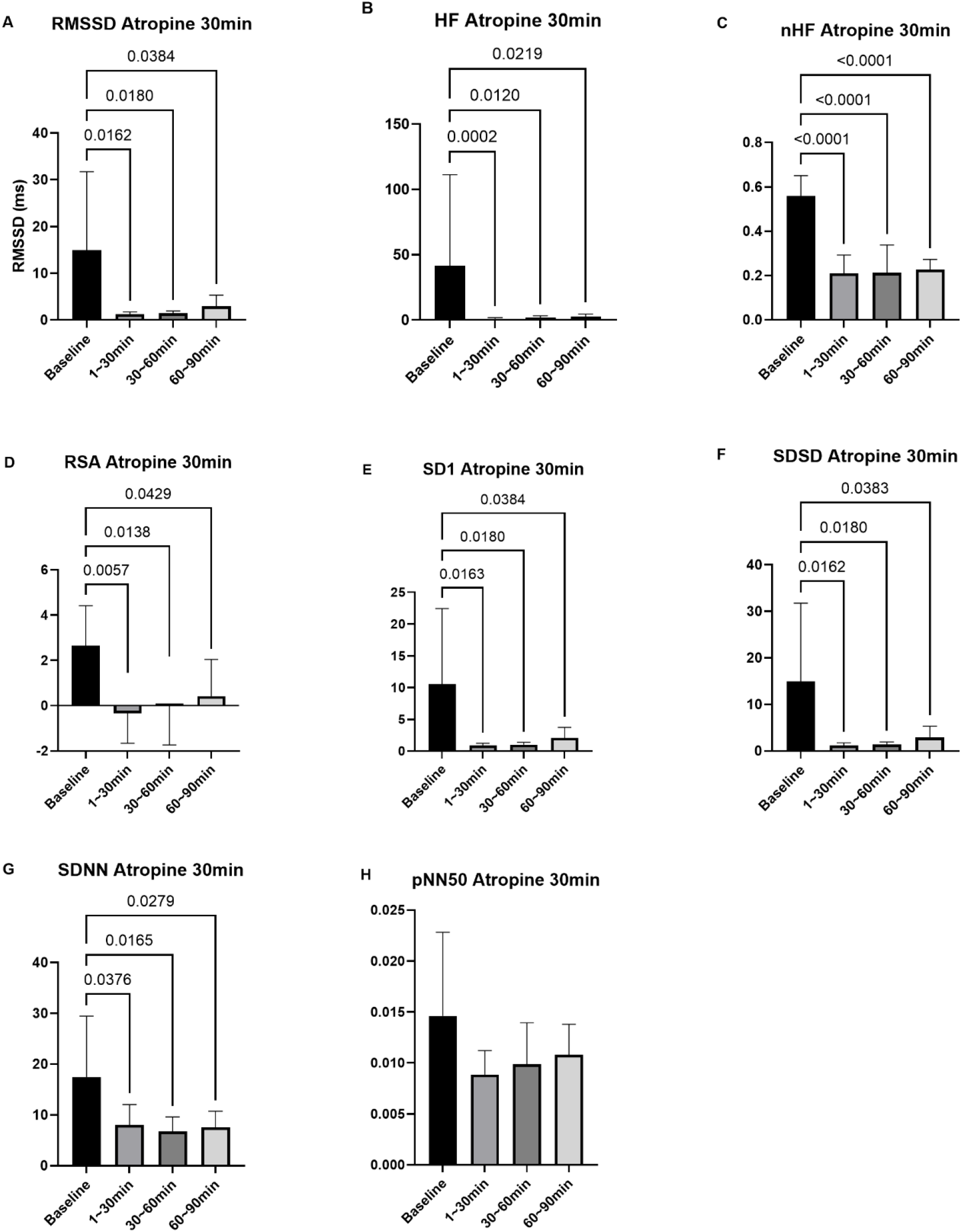
Comparison of parasympathetic-related HRV parameter values in 30-min segments after intraperitoneal atropine administration in rats. HRV parameters were analyzed during baseline and three post-atropine 30-min segments: 1–30, 30–60, and 60–90 min. Panels show: (A) RMSSD, (B) HF Power, (C) HFn, (D) RSA, (E) SD1, (F) SDSD, (G) SDNN, and (H) pNN50%. Data are from 7 rats.

**Figure 9:**
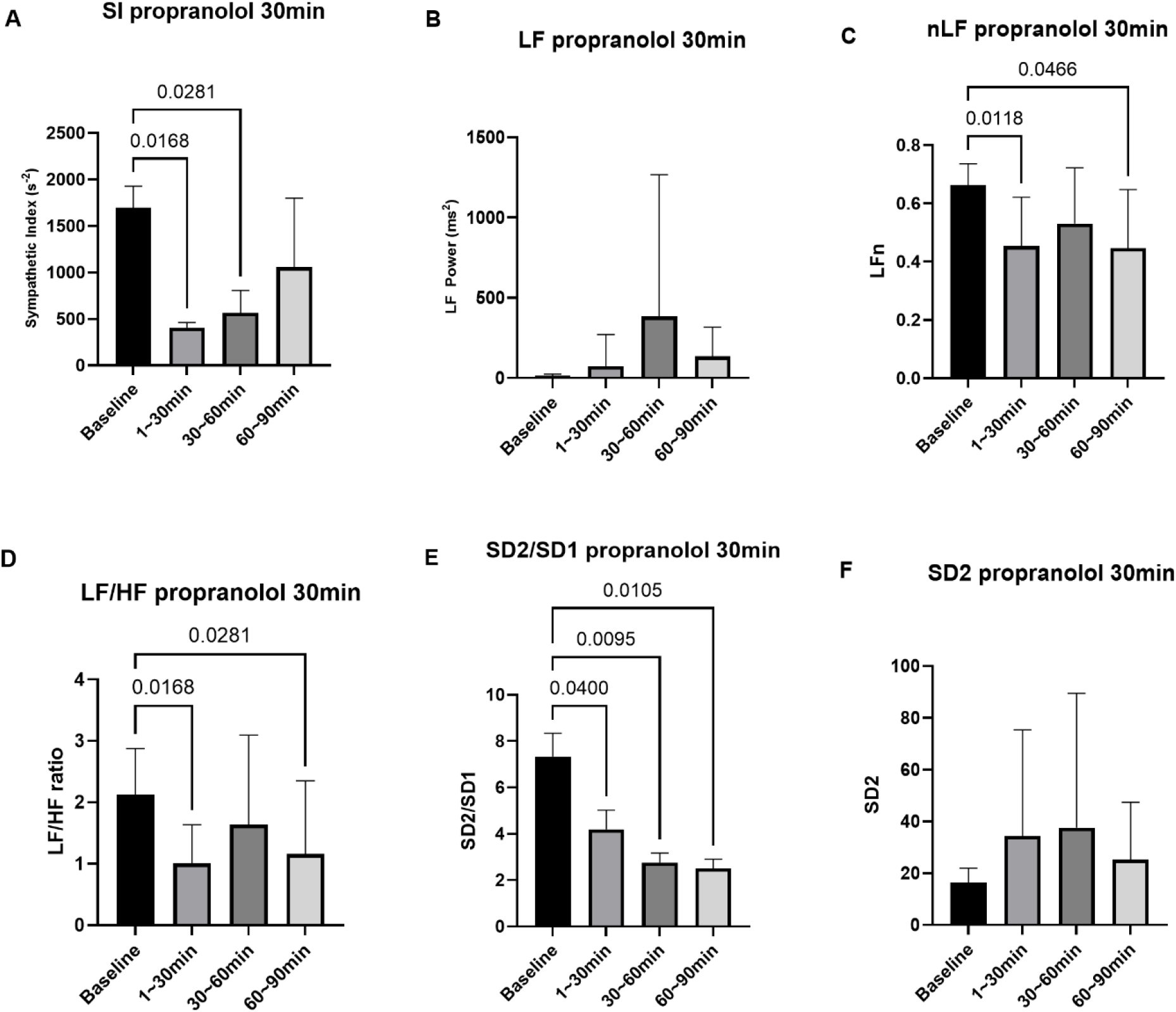
Comparison of sympathetic-related and sympathovagal-balance HRV parameter values in 30-min segments after intraperitoneal propranolol administration in rats. HRV parameters were analyzed during baseline and three post-propranolol 30-min segments: 1–30, 30–60, and 60–90 min. Panels show: (A) Sympathetic Index, (B) LF Power, (C) LFn, (D) LF/HF ratio, (E) SD2/SD1 ratio, and (F) SD2. Data are from 10 rats.

#### Sympathetic-related and sympathovagal-balance HRV responses in 30-min segments after propranolol administration

In the 30-min segment analysis after propranolol administration, Sympathetic Index (SI) detected β-adrenergic sympathetic blockade during the first two post-drug segments. SI was significantly reduced compared with baseline during the 1–30 min and 30–60 min segments, but the effect was no longer significant during the 60–90 min segment, yielding an average p-value of 0.0393 and a sustainability score of 2 out of 3. LF Power did not show significant changes in any 30-min segment. LFn decreased significantly during the 1–30 min and 60–90 min segments but not during the 30–60 min segment, indicating a less consistent response than SI. Similarly, LF/HF ratio was significantly reduced during the 1–30 min and 60–90 min segments but not during the 30–60 min segment. In contrast, the sympathovagal-balance parameter SD2/SD1 was significantly reduced across all three 30-min post-propranolol segments, with the lowest average p-value of 0.0200 and the highest sustainability score of 3. Because SD2/SD1 reflects autonomic balance rather than isolated sympathetic activity, it was interpreted separately from SI. SD2 did not show significant changes after propranolol administration.

## Discussions

This study provides an in-depth analysis about the reliability, sensitivity, and sustainability of the HRV parameters via selective autonomic blockade using propranolol and atropine in rat models. Despite the wide use of heart rate variability parameters for the non-invasive assessment of the autonomic nervous system and development of several HRV parameters, there remains an ongoing debate regarding the optimal parameters for assessing sympathetic and parasympathetic activity. This study addresses this debate by evaluating and ranking HRV parameters for their effectiveness in assessing parasympathetic and sympathetic activities.

Atropine, a muscarinic antagonist, known to significantly reduce the parasympathetic activity (McLendon and Preuss 2024), was used to test the parasympathetic HRV parameters. Atropine is known to act within 10 to 15 minutes after the IM administration, with a duration of action of 90 minutes (“Intramuscular Sedation” 2018). The atropine suppressed parasympathetic activity for all the animals irrespective of their baseline HRV activity, which is represented by the higher standard deviation of the baseline HRV parameters compared to that after atropine suppression. For example, the standard deviation of RMSSD for all the rats during the baseline was 18.72ms which was reduced to 0.6ms in the first 15-min after atropine injection. Similarly, the standard deviation of HF Power was 9.50 in baseline, and it was reduced to 0.62ms^2^ in the first 15 minutes and remained less than 3 ms^2^ upto 75 minutes. The parasympathetic blockade via atropine was better detected by the frequency domain parameters HF power and RSA which are also derived from the value of HF Power. Both the HF power and RSA detected parasympathetic suppression, starting immediately after the administration of atropine up to 60 minutes. Both had the highest sustainability score of 4, compared to other parameters, but based on the average of p-values, the HF Power was slightly better than the RSA. Time domain parameters on the other hand, could not detect the PNS suppression in first 15-min segment, RMSSD and SDSD, and nonlinear parameter SD1, indicated similar behavior, with same average of their p-values, and same sustainability score. They captured the parasympathetic suppression for 15-30 minutes and 30-45 minutes segments only. SD1 have been known to be similar to RMSSD (Shaffer and Ginsberg 2017), which is reconfirm by our results.. PNN50% also detected PNS suppression from 15-60 minutes but could not capture it in the first 15 minutes, hence its sustainability score was less than HF Power. SDNN was not influenced by atropine administration. Normalized HF Power (HFn) showed moderate reliability in the 15-min analysis, detecting parasympathetic blockade during the 15–30 and 30–45 min segments but not during the first 15-min segment. However, when HRV was analyzed using 30-min windows, HFn became the most statistically reliable parasympathetic-related parameter, with the lowest average p-value and a maximum sustainability score. This suggests that HFn may require a longer recording window to reliably detect atropine-induced parasympathetic suppression.

To study the effect of signal duration on HRV parameter reliability, HRV parameters were recalculated using 30-min post-drug segments instead of 15-min segments. Using 30-min windows, RMSSD, SDSD, SDNN, and SD1 detected parasympathetic suppression within the first 30-min segment and remained significantly lower than baseline up to 90 min, similar to HF Power, RSA, and HFn. These findings suggest that time-domain and nonlinear Poincaré plot parameters may require longer signal durations than frequency-domain parameters to reliably detect rapid autonomic changes.

Propranolol, a non-selective β-adrenergic blocker, is known to significantly reduce the sympathetic activity (Bylund 2015; Ammar and Hussein 2018; Shahrokhi and Gupta 2024), was used to suppress the sympathetic cardiac output, to study if the HRV variables used to assess the sympathetic nervous system activity can capture this change and to rank them on the basis of their sensitivity and sustainability. Propranolol is known to start acting within 30-60 min and its effect lasts for 3-4 hours (PAterson et al. 1970; Coltart and Shand 1970).

The results indicated that the Sympathetic Index (SI), captured the suppression of sympathetic activity after 15 minutes of drug administration, with the sustainability score of 2, compared to the LF power and normalized LF power (LFn). Low Frequency band of the power spectrum of the HRV signal is known to be influenced by both the sympathetic as well as parasympathetic activity (Ali and Chen 2023), but it has been widely used as measure of sympathetic activity. Our results reaffirm that LF should not be used as an isolated indicator of sympathetic activity. While the normalized LF power detected the initial sympathetic blockade in the first 15 minutes, it could not detect this change after 15 minutes. SD2 was unaffected by propranolol administration. The parameters of autonomic balance, LF/HF also detected the initial change in first 15 minutes but could not detect afterwards, while SD2/SD1 was not affected at all.

Increasing the time segment used to calculate the HRV variables from 15-min to 30-min post drug administration, the SI was significantly lower than the baseline value within the first and 2nd 30-min segments (up to 60 min) before it started increasing. The frequency domain parameters LF power was not affected at all, while LFn showed decrease within first and 3^rd^ 30-min segment post propranolol administration, but not during the 2^nd^ segment (30-60 min), which is similar to LF/HF ratio. However, the autonomic balance HRV parameter SD2/SD1 (which is derived from the Poincare plot) indicated the shift in the autonomic balance towards parasympathetic nervous system during all the 30-min time segments. Which indicated that the Poincare plot is more reliable with increased signal duration (30-min instead of 15-min).

Based on our results, HF Power was the most reliable frequency-domain parameter for detecting rapid parasympathetic suppression when shorter 15-min HRV analysis windows were used. In contrast, HFn showed improved reliability when HRV was analyzed using longer 30-min windows, suggesting that normalized frequency-domain measures may require longer recording durations to more consistently capture atropine-induced parasympathetic blockade. For sympathetic-related assessment, LF Power was not reliable, likely because LF Power reflects mixed sympathetic and parasympathetic influences rather than isolated sympathetic activity. SI showed the most reliable sympathetic-related response after propranolol administration, whereas SD2/SD1 showed a strong sympathovagal-balance response in 30-min segments. Overall, these findings indicate that HRV parameter selection should account for both the autonomic branch being evaluated and the analysis-window duration. This approach provides a structured framework for selecting appropriate HRV measures in experimental autonomic function studies.

## Ethics Statement

All animal procedures were approved by the Institutional Animal Care and Use Committee of the University of Michigan and were conducted in accordance with institutional guidelines for the care and use of laboratory animals.

## Funding

This study was partially supported by NIH grant R01DK131524

## Conflict of Interest

The authors declare no competing interests.

## Author Contributions

J.D.Z.C. conceived, designed, and supervised the study, provided scientific guidance, and reviewed and edited the manuscript. M.K.A. performed the electrode implantation surgeries, animal experiments, animal care, ECG recordings, HRV data analysis, figure preparation, and drafted the manuscript. S.G. assisted with data analysis. S.J. and S.L. assisted with animal care, surgical procedures, and drug administration. All authors reviewed and approved the final manuscript.

## Notes

### Competing Interest Statement

The authors have declared no competing interest.

